# Identification of novel Bromodomain inhibitors of *Trypanosoma cruzi* Bromodomain Factor 2 (*Tc*BDF2) using a fluorescence polarization-base high-throughput assay

**DOI:** 10.1101/2024.02.16.580721

**Authors:** Luis E. Tavernelli, Victoria L. Alonso, Imanol Peña, Elvio Rodríguez Araya, Romina Manarin, Juan Cantizani, Julio Martin, Juan Salamanca, Paul Bamborough, Felix Calderón, Raquel Gabarro, Esteban Serra

## Abstract

Bromodomains are structural folds present in all eukaryotic cells that bind to other proteins recognizing acetylated lysines. Most proteins with bromodomains are part of nuclear complexes that interact with acetylated histone residues and participate in regulating DNA replication, transcription, and repair through chromatin structure remodeling. Bromodomain inhibitors are small molecules that bind to the hydrophobic pocket of bromodomains, interfering with the interaction with acetylated histones. Using a fluorescent probe, we have developed an assay to select inhibitors of the bromodomain factor 2 of *Trypanosoma cruzi* (*Tc*BDF2) using fluorescence polarization. Initially, a library of 28,251 compounds was screened in an endpoint assay. The top 350 ranked compounds were further analyzed in a dose-response assay. From this analysis, 7 compounds were obtained that had not been previously characterized as bromodomain inhibitors. Although these compounds did not exhibit significative trypanocidal activity, all showed *bona fide* interaction with *Tc*BDF2 with dissociation constants between 1 and 3 μM validating these assays to search for bromodomain inhibitors.

## Introduction

*Trypanosoma cruzi* is a unicellular parasite that causes Chagas Disease (also known as American trypanosomiasis). It is estimated that at least 6 million people are infected globally, and around 12,000 deaths occur annually because of this illness. In the Americas, Chagas disease is the most important parasitic disease, with an annual incidence of 30,000 new cases on average, among them 8,600 are newborns that become infected during pregnancy ^1^. Originally, Chagas disease was constrained within Latin American countries, but recent emigration movements have spread the infection to other territories such as Spain, the USA, or Canada ^2^. Only two active compounds are currently being used to treat the disease: Nifurtimox and Benznidazole, both developed more than fifty years ago. These compounds display severe side effects and even though their use in the acute phase of the disease is efficient, their use in the chronic phase is still controversial ^3^. These facts highlight the need for new, improved, and efficient drugs against this deadly disease.

Lysine acetylation is a reversible post-translational modification (PTM) found in a myriad of proteins, but the exact function of this PTM is just starting to be understood. In the past years, the target proteins that were subjected to this modification were found to be involved in distinct nuclear processes such as transcription, DNA replication, and repair, presumable due to chromatin remodeling ^4^. Nowadays, acetylation is known to be present in all cellular compartments participating in diverse processes like energetic metabolism, protein degradation, protein localization, and cell cycle regulation, among others ^5,6^.

Bromodomains (BD) are ∼110 amino acid-long protein modules that specifically recognize and bind to acetylated lysines (AcK). These domains bear a left-handed four-α-helix bundle structure (αA, αB, αC, and αZ) connected by two loops (ZA and BZ loops) that form the accessible hydrophobic pocket where the recognition of the Ac-K takes place ^7^. Currently, it is known that there are eight coding sequences for BD-containing proteins in trypanosomatids, named *Tc*BDF1 to *Tc*BDF8 ^8,9^. Our lab has been pioneering in the characterization of BDs in *T. cruzi* throughout the years ^10–14^. The first bromodomain to be studied in our group was *Tc*BDF2, which is expressed throughout the parasite’s life cycle and is found in discrete regions within the nucleus ^12,14^. Furthermore, this protein bears a bipartite Nuclear Localization Sequence (NLS) that targets the bromodomain in the nucleus. Also, *Tc*BDF2 was shown to be important in the devolvement of infection *in vitro* as well as the kinetic of the replicative stages in epimastigotes. We have shown that this bromodomain can interact with histone H2 and H4 (through the acetylated lysines 10 and 14). In addition, a recent study demonstrated that *Tc*BDF2 was associated with H2B.V and directly interacts with the histones H2B.V, H2B, and H4 in *in vitro* assays ^15^.

Out of a limited number of cases with cytoplasmic or dual localization ^10,11,16–18^, BDs compose a family of proteins that are usually known as “readers” that recognize acetylation marks on histones. Once the “reading” takes place, bromodomains act as bridges or scaffolds for the assembly and/or recruitment of other factors allowing the interaction with the chromatin within that region ^19^. BD-driven recruitment of complexes into the chromatin influences key regulatory processes within the nucleus such as transcription, DNA repair, and DNA replication, among others. All these processes must be carried out in a precise and regulated manner, where an unbalanced regulation or incorrect function of BD factors can be associated with cell death or uncontrolled proliferation. For example, in many types of cancers or chronic inflammatory diseases, BD factors are upregulated and considered potential targets for new drug discovery campaigns ^20^.

The development and discovery of small molecules that can bind and inhibit proteins is a rapidly advancing field of research. In the past years, many molecules against bromodomains have been identified, proving their efficacy as active inhibitors. To date, according to the Clinical Trials Database from the National Academy of Medicine, there are 55 studies (either completed or still running) with BD inhibitors mainly targeted to different types of cancer and inflammatory diseases, among others (https://clinicaltrials.gov/, last consultation 12/12/23). In this context, parasitic bromodomains have also been gaining attention as attractive targets to battle NTDs such as American trypanosomiasis, Leishmaniasis, Malaria, and Toxoplasmosis ^21–27^.

In a previous report, we established the relevance of *Tc*BDF2 in all the stages of the parasite and described a small set of molecules, already known as mammalian BD inhibitors that were able to bind specifically to *Tc*BDF2 ^14^. In the present study, we describe a high-throughput competition assay based on Fluorescent Polarization (FP) that allowed us to identify molecules that bind to *Tc*BDF2. After this initial screening, the binding of the selected compounds was confirmed by Thermal shift and the cytotoxicity was tested against different life cycle stages of *T. cruzi*. We were able to identify *Tc*BDF2 inhibitors that serve as a starting point for rationally designing new compounds to be explored against Chagas disease.

## Material and Methods

### Protein Purification

*Escherichia coli BL21* carrying pDEST17-*Tc*BD2 and pDEST17-*Tc*BD2m (mutant BD2: Y85A and W92A without the ability to bind to acetylated Histone 4^28^) were grown at OD ∼ 0,6 and induced with 0.1 mM isopropyl-b-D-thiogalactopyranoside during 4 hours at 37°C as described previously ^14^. The proteins were purified using Ni-NTA (Thermo Fisher™) following the manufacturer’s instructions. The purified proteins were dialyzed against the previously determined optimal assay buffer: phosphate 0.1 M PH=8, glicerol 1%, DMSO 0.5%, and diluted to a final concentration of 200 μM. Correct secondary structure of soluble proteins was verified by circular dichroism spectroscopy using a spectropolarimeter (Jasco J-810, Easton, MD, USA).

### Fluorescence polarization (FP) assay setup and data analysis

FP measurement was performed on a BMG PheraStar FS plate reader (BMG Labtech GmbH, Ortenberg, Germany) at an excitation wavelength of 488 nm and an emission wavelength of 675 nm (50 nm bandwidth) and Black 1536-well flat bottom small volume microplates with a non-binding surface from Greiner Bio-One GmbH (Frickenhausen, Germany). Optimal Bromosporine (BSP) probe/*Tc*BD2 interaction conditions were assayed by including in the reaction phosphate medium and seriated dilutions of DMSO (up to 2%), DTT (up to 50 μg/ml), EDTA (up to 90 μg/ml), glycerol (up to 10%), BSA (up to 250 μg/ml), deoxy big-chaps (up to 50 μg/ml), CHAPS (up to 50 μg/ml), Zwittergent 3-14 (up to 50 μg/ml), Triton-X100 (up to 0,5 M), NP40 (up to 0.3 M), Tween-20 (up to 100 μg/ml) and Pluronic acid (up to 10%). The final conditions of the screening assay were set at a final volume of 8 μl, 50 nM of the Bromosporine probe (BSP-AF488), and 100 μM of *Tc*BDF2, phosphate pH=8, glycerol 1 %, and DMSO 0.5 %. Recombinant *Tc*BD2m was used as a negative control of binding. For the screening, 8 microliters of recombinant *Tc*BD2 contained in the assay buffer plus 50 nM of BSP-AF488 were dispensed in 1,536 wells Grenier plates were compounds resuspended in 8 μL of DMSO, were previously dispensed. The primary screening was performed at a single shot at a final concentration of 100 μM *per* well. Further on, to determine the compounds’ potency 11-concentration points in a 1:3 seriated dilutions pattern were stamped in the plate, starting at 100 μM. Plates were stored frozen at -20°C until used when they were allowed to equilibrate at room temperature before proceeding to the reading. Controls were made by using recombinant *Tc*BD2m instead of *Tc*BD2, under the same conditions. All manipulations were made by using Multidrop™ Combi Reagent Dispenser.

Statistics, Z values, and robustness (3SD) to determine activate cutoff were calculated using templates in ActivityBase.

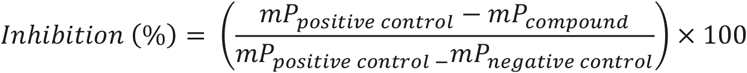

The validation of the assay performance for each enzyme was quantified by calculation of the Z′-factor using the formula:

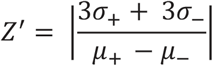

where σ+ and σ− are the standard deviations and μ+ and μ− are the mean values of the positive and negative controls, respectively. A series of negative and positive controls was measured. For each, positive and negative control, 64 wells were analyzed.

### Thermal shift

Recombinant *Tc*BD2 was buffered in 10 mM HEPES, pH 7.5, and 300 mM NaCl and assayed in a 96-well plate at a final concentration of 5 μM in a 25 μL volume. Compounds were added at a final concentration between 1 and 50 μM to calculate dissociation constants. SYPRO Orange was added as a fluorescence probe at a dilution of 1:10000. Excitation and emission filters for the SYPRO Orange dye were set to 465 and 590 nm, respectively. The temperature was raised with a step of 2°C per minute from 25°C to 96 °C, and fluorescence readings were taken at each interval in a real-time Biorad Opus CFX. Data were analyzed as previously described ^29^.

## Molecular Modeling

### Protein preparation and grid generation for docking

The crystal structure of *Tc*BDF2 solved with bromosporine (PDB entry 6NIM, resolution = 1.78 Å) was used as a reference structure for modeling in Maestro (Schrodinger Release 2022-2: Maestro Schrodinger LLC: New York, 2022) ^30^. The selection of the structure was based on the occupation of the bromosporine at the active site and the good resolution. The Protein Preparation Wizard module was used to remove solvent molecules, add missing side chains (Thr9, Lys27, Lys45, Lys64, Lys88 placed outside of the binding pocket), add hydrogens (PROPKA pH 7.0), and minimize the structure under the OPLS4 force field with heavy atoms restrained to 0.3 A RMSD. Additionally, four buried water molecules (W 304, 305, 316, 327 according to the residue number of PDB 6NIM; see SI) were retained at the bottom of the binding cavity 42-43. The grid was defined as a closed box centered at the bromosporine, and other settings were set to default values.

### Ligand preparation for docking

Chemical structures (compounds) were prepared using LigPrep to generate the 3D conformations. The protonation states were generated at pH 7.4 ± 0.5 and geometry optimization with the S-OPLS force-field.

### Docking protocol

Docking was performed using Maestro with the extra precision (XP) mode using Glide. Ligand sampling was set to flexible and Epik state penalties to docking score were included. The hydrogen bonds with Asn86 and Trp304 were selected as constraints where at least one of them must match.

### Beta-galactosidase expressing parasites assay

*T. cruzi* Dm28*c* which expresses the *Escherichia coli* LacZ gene was used ^31–33^. Epimastigotes were incubated with the compounds (50, 25, 12.5, 6.25 and 3.125 μM). After 72 h of treatment the assays were developed by the addition of Chlorophenol red-β-D-galactopyranoside (CPRG) (100 μM final concentration) and Nonidet P-40 (0.1% final concentration). Plates were incubated for 2 to 4 hours at 37°C. Wells with β-galactosidase activity turned the media from yellow to red, and this was quantitated by Absorbance at 595 nm using a Synergy HTX multi-detection microplate reader as reported previously^31^. Normalized survival percentage was plotted against concentrations on Prism 9.0 GraphPad software. DMSO 50% was used as the zero percent survival baseline. Each concentration was assayed in triplicates.

### MTT Assay

Cell viability after treatment was determined by the 3-(4,5-dimethylthiazol-2-yl)-2,5-diphenyltretazolium bromide (MTT) reduction assay as previously described ^29^. Briefly, Vero cells (5.000 cells per well) were incubated in a 96-well plate in the presence of each compound (200, 100, 50, 25 and 12.5 μM) for 48 hours. Then 200 μL MTT solution (5 μg mL-1 in PBS) was added to each well and incubated for 1 hour at 37°C. After this incubation period, the MTT solution was removed and precipitated formazan was solubilized in 100 μL of DMSO. Optical density (OD) was spectrophotometrically quantified (λ= 540 nm) using a Synergy HT multi-detection microplate reader. DMSO was used as blank, and each treatment was performed in triplicates. IC_50_ values were obtained using non-linear regression on Prism 9.0 GraphPad software.

## Results and discussion

### Identification of *Tc*BD2 binders through a fluorescence polarization-based high throughput screening

To identify molecules that bind to the *Tc*BDF2 bromodomain (*Tc*BD2), we set up a high-throughput screening (HTS) assay using Fluorescent Polarization (FP). FP is a very sensitive technique that allows quantitative analysis of the interaction between two molecules. Recent developments have allowed this technique to be adapted to be used in HTS format toward finding inhibitors against a myriad of proteins ^34,35^.

Firstly, we assayed previously used fluorescent probes with an affinity for the mammalian BDs ATAD2^36^, PCAF and GCN5^37^, BRPFs^38^, Brd9 that is also a promiscuous human Brd binder^39^, BET BD1 selective^40^ and BET BD2 selective (Patent WO2014140076). None of these probes showed significant fluorescence polarization when incubated with recombinant *Tc*BD2 under standard conditions, reinforcing the idea that these parasite BDs are very divergent from human ones.

Given that we previously showed that the BD-pan inhibitor Bromosporine (BSP) binds to *Tc*BD2 ^14^, we decided to modify this molecule to use it as a fluorescent probe in the HTS assay. Hence, AlexaFluor-488 (AF488) was coupled to BSP (Fig. S1). BSP was our last choice because we have previously demonstrated that it has no significative trypanocidal activity ^13^. BSP-AF488 probe was used with the wild-type (*Tc*BD2) and a previously characterized recombinant mutant version of *Tc*BD2 (*Tc*BD2m, unable to bind to its acetylated ligand) to optimize the conditions in a HTS format. We first tested different buffer compositions to obtain a working window of approximately 120 mP, needed to proceed to the screening assay. Optimal conditions for the screening were determined as shown in Figure 1. Screenings were made in a 1,536 plate with a final volume of 8 μL. The condition selected was: 100 μM of recombinant *Tc*BD2, 50 mM of BSP-AF488, buffer phosphate 0.1 M, glycerol 1 %, and DMSO 0.5 %. Recombinant *Tc*BD2m was used as a negative control in all plates assayed. As can be observed, there is very low interaction between the probe and BD2m in all conditions tested. The Z’ score, calculated as mentioned in the Methods section, was used as a quality control parameter ^41^.

**Figure 1:**
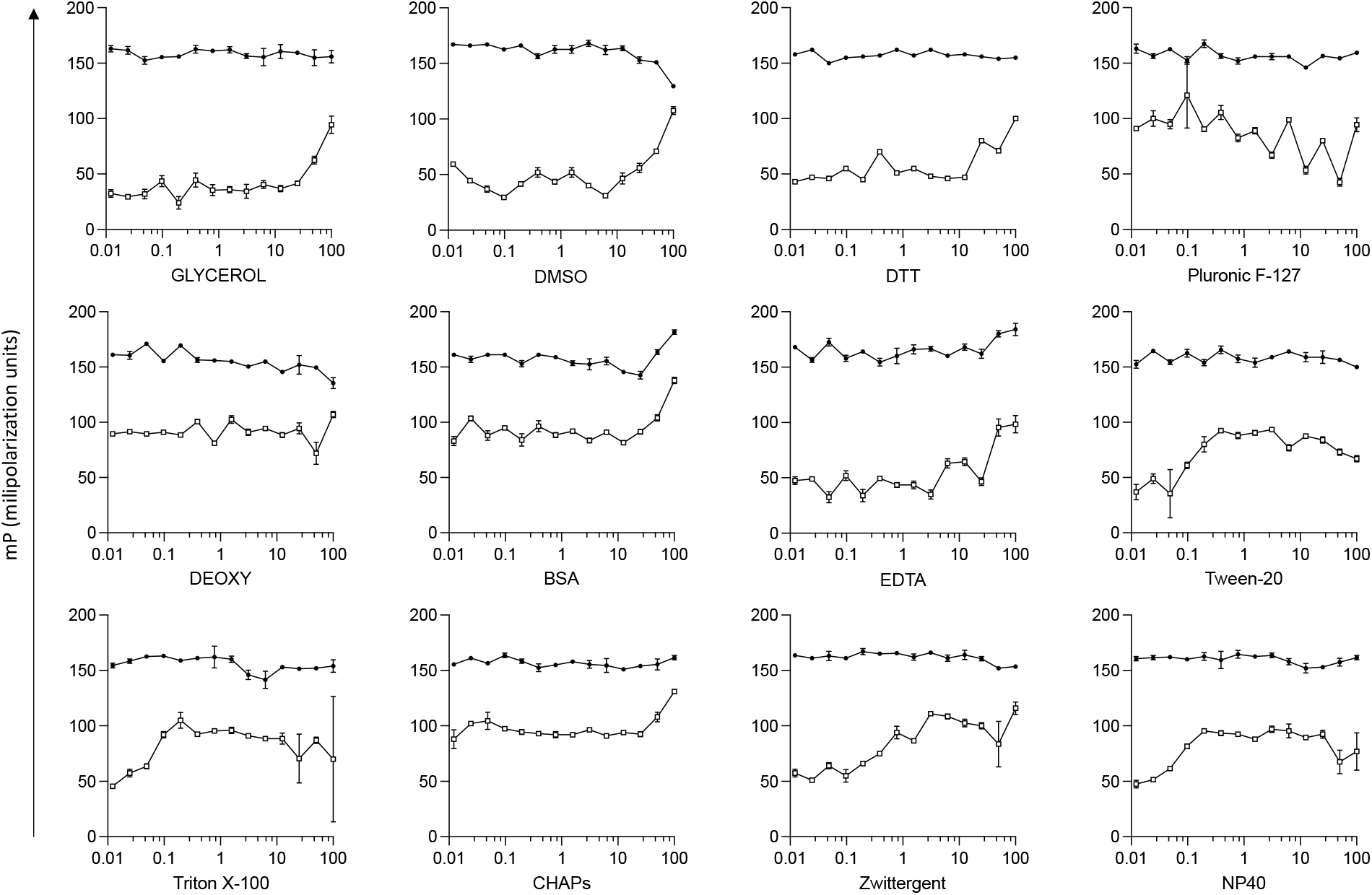
Fluorescent polarization assay setup. Different additives to the buffer used for the assay were tested to find the best conditions. All indicated reagents (below the graphs) were assayed in a 50 mM phosphate buffer pH=8. Concentrations assayed are indicated in the Methods section and were normalized in a logarithmic scale on the X-axis. On the y-axis polarization fluorescence is indicated in mP units. Recombinant *Tc*BD2m was used as a negative control of binding. Grainer plates of 1536 wells were used for all assays and the reading was made with PheraStar Technology. (O)TcBD2m, TcBD2WT (•).

Once the assay conditions were established, we performed the screening of a small molecule bromodomain-targeted compound set (summarized in Fig. 2A). The set mostly consisted of compounds showing experimental activity against bromodomains or synthesized during various human bromodomain drug discovery programs, and their analogs (total of 28,251 compounds). The library was tested in a single shot format at a final concentration of 100 μM, in a 1,536-wells plate format. All plates assayed were analyzed using in-house algorithms and the Z’ score (over 0.4) was used as a quality control parameter.

**Figure 2:**
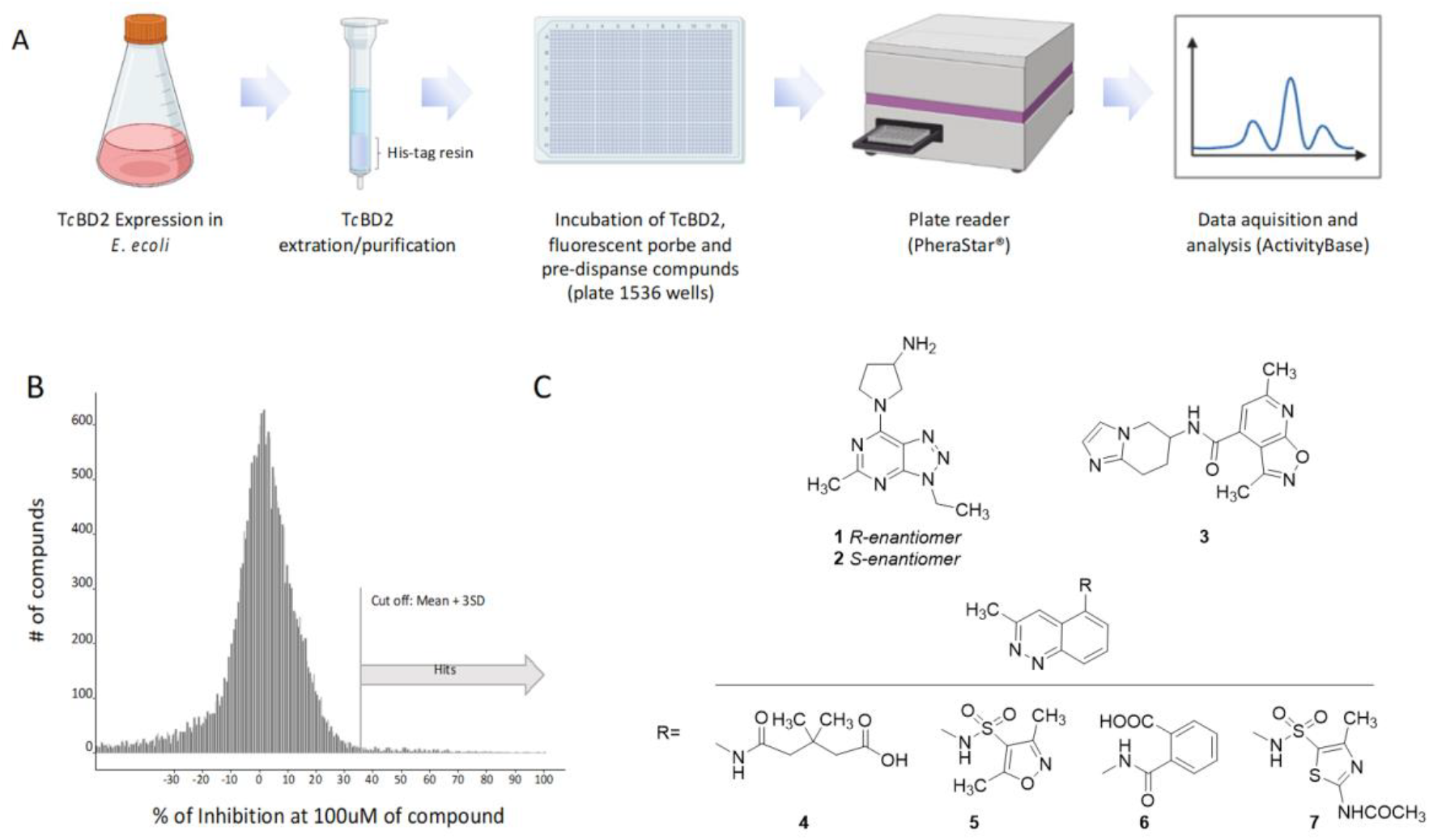
HTS assay to identify TcBD2 binders. A) Schematic flowchart of HTs assay. B) Histogram representing 28,251 compounds assayed in a single shot format using 1,538 grainer plates. All compounds were tested at a final concentration of 100 mM. The cut-off to consider positive hits was set at 37% of inhibition. *Tc*BD2 was used as positive (100% fluorescent probe bound to the BD) and *Tc*BD2m as negative (100% fluorescent probe non-bound to BD). Z’ score was used as the quality control parameter where only plates with Z’ score of above 0.4 were considered for analysis. C) Structures of hits with pIC50 ≥ 4,5 resulting from the seriated dilutions screening of the 350 selected compounds (all compouds were >90% purity). Figure 2A was created with Biorender.com.

Out of all compounds tested, 471 compounds were considered positive after the first single shot format assay was performed. The cut-off for positive hits was set at (mean + 3SD) or 37% inhibition, having a positive hit rate of 1.7% (Fig. 2B). Next, to determine the compounds’ potency, 11 concentration points in a 1:3 dilution pattern were stamped in a 1,536-plate starting at 100 μM. Recombinant *Tc*BD2m was used as a negative control. In this assay, 7 compounds showed binding activity of pIC50 ≥ 4,5 against *Tc*BD2 in a dose-response manner (Fig. 2C, Table 1).

**Table 1.** Dissociation constants and binding measured by thermal shift over recombinant *Tc*BD2.

| Compound | 1 | 2 | 3 | 4 | 5 | 6 | 7 |
| --- | --- | --- | --- | --- | --- | --- | --- |
| *pKd* | 5.8 | 1.9 | 5.6 | 5.8 | 3.52 | 5.4 | 5.5 |
| *pIC50* | 4.6 | 4.5 | 4.7 | 5.1 | 5.6 | 4.5 | 4.8 |

The binding of the 7 selected compounds to *Tc*BD2 was verified by differential scanning fluorimetry (DSF) or Thermal shift observing *Kd*s between 1 and 3 µM (Table 1 and Fig. S2). The (most promising) compounds were modeled at the AcK site of *Tc*BDF2 to further validate their potential role as inhibitors (Fig. S46). Importantly, AcK binds in a heavily hydrophobic pocket but contains several well-resolved structural waters in the binding pocket. These water molecules are observed at the base of the pocket in most BRD as well as in the X-ray structure of *Tc*BDF2 (Fig S3) ^42,43^. AcK recognition is mediated by a direct hydrogen bond to a conserved asparagine (Asn 86 in *Tc*BDF2), which is mimicked by most BRD inhibitors. Moreover, there is also a conserved tyrosine (Tyr43 in *Tc*BDF2) to coordinate the active site bridging water molecule^44^ (Fig. S3). Both interactions are also conserved in the crystal structure of *Tc*BDF2 with BSP. We evaluated whether the compounds were able to retrieve those interactions at the AcK pocket through molecular modeling (while binding to the AcK pocket). The docking procedure was validated by reproducing the binding mode of bromosporine while maintaining key interactions (Fig. S4). Docking studies revealed that all seven compounds fit into the substrate pocket of *Tc*BDF2. The compounds were able to form a hydrogen bond with Asn86 and/or the water-bridged hydrogen bond with Tyr43 (Fig. S5-S6). It is worth mentioning that compounds 4, 5, 6, and 7 with methylcinnoline as core made pi-stacking interaction with Trp92, and even 4 and 5 can form a hydrogen bond in a similar way to BSP (Fig. S5-6). The docking pose of compound 3 placed the dimethyl-isoxazole-pyridine moiety rotated 180º vertically from the expected position, as is the case with the AcK mimic methylbenzoisoxazole (PDBid 5Y8C)^45^. The enantiomer compounds 1 and 2 also made a pi-cation interaction with Trp92, where they can also form pi-stacking interactions between the 3-ethyl-5-methyl-triazolopyrimidine core and Trp92 (Fig. S6).

Finally, the 7 compounds were tested in an *in vitro* assay of *T. cruzi* intracellular amastigote assay using VERO cells as hosts ^46^ and in an *in-house* colorimetric assay using epimastigotes of the Dm28c strain that expresses beta-galactosidase from an episomal plasmid (Fig. S7)^31^. Not surprisingly, none of the compounds showed parasiticidal activity at concentrations below 50 μM in the two life cycle stages. Additionally, we assessed the toxicity of these compounds in the cell line used for the infection assays using MTT. Our results indicate that none of the compounds exhibited cytotoxicity up to 200 μM (data not shown).

## Conclusions

We have previously assayed the binding of *Tc*BDF2 and *Tc*BDF3 to several BD inhibitors and determined that they have different binding specificities ^13,14^. Commercial human inhibitors iBET-151 and BSP bind to both bromodomains; but JQ1(+), which binds *Tc*BDF3 with an affinity similar to iBET-151, does not interact with *Tc*BDF2. The compounds from the HTS we tested herein have *Kd*s for *Tc*BD2 between 1 and 3 μM, however, they have no significative activity against epimastigotes, nor amastigotes in infected cells. This effect could be due, at least in part, to a low potency of the compounds obtained in our screening, a fact that could be associated with the use of BSP as a probe, a compound that, itself, shows low trypanocidal activity. Also, a possible explanation for this lack of activity of *Tc*BDF2 inhibitors could be associated with the multimeric nature of the complexes in which bromodomains are included (see below).

The human inhibitors mentioned were also assayed against BDF orthologues from other trypanosomatids with different results. Schulz and coworkers showed that iBET-151 (BET bromodomains inhibitor) induces *T. brucei* bloodstream form to develop insect-stage features, like the expression of surface procyclin and upregulation of glycolysis enzymes. They found that *Tb*BDF2 and *Tb*BDF3 bind iBET-151 with a low affinity (Kd=225 μM and 175 μM, respectively) and do not bind at all to other BET inhibitors like JQ1(+). iBET-151 was co-crystalized with *Tb*BDF2, and it was found in a completely atypical position, flipped by roughly 180° to the position it binds to human BDs, something that could explain the low affinity of the interaction ^21,47^. Later, Yang and collaborators assessed 27 compounds obtained by a structure-based virtual screening combined with ITC experiments. They found one compound (GSK2801) that binds with higher affinity to the BD of *Tb*BDF2 (Kd=15 μM) and with lower affinity to the second BD of *Tb*BDF5 (Kd=83 μM). By contrast, GSK2801 does not bind to *Tb*BDF3 or the first BD from *Tb*BDF5 ^48^. BDF5 from *Leishmania donovani* was also recently assayed against human BD inhibitors ^27^. *Ld*BDF5 binds to SGC-CBP30, BSP, and I-BRD9, with different affinities for the first or second DB. SGC-CBP30, which showed the higher affinity (Kd=281 nM, for *Ld*BD5.1), also exhibited activity against promastigotes from *L. mexicana* (IC_50_=7,16 μM) and *L. donovani* (IC_50_=6,16 μM).

It is worth mentioning that not always a direct correlation is found between the affinity of the inhibitors and their activity against the parasite in *T. cruzi* nor *T. brucei or Leishmania*. For example, iBET-151 and JQ1(+) have IC_50_ against Dm28c epimastigotes of 6.35 μM and 7.14 μM respectively ^13^. In contrast, BSP that has a *Kd* for *Tc*BDF2 similar to iBET-151 was less active against parasites (IC_50_ > 50 μM). GSK2801 inhibits the growth of procyclic *T. brucei*, disrupting the nucleolar localization of *Tb*BDF2, with an IC_50_ of 1.37 μM, which suggests that GSK2801 has other targets beyond the inhibition of *Tb*BDF2 ^48^. Also, as was mentioned above, the IC_50_ for SGC-CBP30 against *Leishmania* promastigotes was significantly lower than the *Kd* measured for this compound on *Ld*BD5.1.

Staneva and coworkers established that the majority of *T. brucei* BDFs participate in complexes that include other BDFs ^49^. A similar situation was reported by Jones and coworkers for *Ld*BDF5 ^50^. In this situation, which seems to be characteristic of trypanosomatids, limited inhibition of only one BD of the complex could be countered by the presence of another one or ones. However, over-expression of dominant negative mutants of one of the BDFs would induce disruption of the whole complex and a more deleterious effect over the parasite, as we have previously reported for *Tc*BDF2 ^28^. This model could also explain the results obtained for other BD inhibitors in other trypanosomatids. Moreover, extrapolating when determining the essentiality of BDFs in these parasites should be made with caution.

The activity of bromodomain inhibitors was also determined for other parasites showing intriguing results. Chua and coworkers assayed 42 compounds previously characterized as BD inhibitors for activity against the asexual form of *Plasmodium falciparum*. All compounds were predicted to be correctly placed into the BD of *P. falciparum* histone acetyltransferase *Pf*GCN5 and to interact with the conserved asparagine by a hydrogen bond, but these interactions were not experimentally measured. SGC-CBP30, a selective inhibitor of human CREBBP (CBP) and EP300 bromodomains, showed the highest *in vitro* activity, with an IC_50_ of 3.2 μM and a selectivity index (calculated using a human HEK 293 cell line) of around 7 ^24^. Finally, *Toxoplasma gondii* was assayed with L-Moses, a specific inhibitor for the GCN5-family bromodomains ^26^. L-Moses interferes with the *in vitro* interaction of GCN5b bromodomain with acetylated histone residues and displays potent activity against *Toxoplasma* tachyzoites infecting HFF cells (IC_50_ of ∼0.6 μM).

All this evidence suggests that maybe targeting only one BD is not the best strategy against trypanosomatids. An alternative could be designing pan-inhibitors against BDs present in nuclear protein complexes taking advantage of the divergence in parasitic BDs *vs*. human BDs. However, it is hard to conceive that such kind of compound could be designed. Another alternative could be focusing on BDs with novel localizations outside the nucleus that are druggable, at least in *T. cruzi*, and do not take part in protein complexes with other BDs ^29,51^.

In summary, we identified 7 hits competitive against *Tc*BDF2 in a fluorescence polarization assay and validated their binding by DSF. The confirmed hits originated from a set of compounds targeting human bromodomains, but despite bearing some features reminiscent of previously published bromodomain inhibitors they are structurally distinct. Except compound 3, all have been tested in various historical hBRD4 assays at GlaxoSmithKline and did not give fitted BRD4 dose-response curves at concentrations up to 50 µM. We are therefore confident that the hits are not generally promiscuous bromodomain inhibitors but may represent specific binders to *Tc*BDF2 with some selectivity over the BET family, the most studied human bromodomain-containing proteins and those whose potential clinical safety risks are best understood. While these hits bind more weakly to *Tc*BDF2 than clinical inhibitors of human BET bind to their targets *in vitro* (typically at least 100 nM), they represent good starting points for optimization.

## Acknowledgments

This work was supported by Tres Cantos Open Lab Foundation (Tc261) and Agencia Nacional de Ciencia y Tecnologıa from Argentina (PICT-2017-1978, PICT-2020-SERIEA-01704) and Universidad Nacional de Rosario (1BIO490). We would like to thank Dolores Campos for their technical assistance with Vero cell and parasite culture.

